# A lightweight codon-based DNA Transformer for Regulatory Region Identification in the Genome

**DOI:** 10.64898/2026.05.04.722647

**Authors:** Ammu Sreenivas Pavan Karthik, Asim Bikas Das

## Abstract

We developed a lightweight codon-based DNA Transformer equipped with multi-head self-attention and an adaptive classifier head, which achieves exon intron classification with high accuracy and also has moderate accuracy in CDS classification and splice site recognition. We named this model as ExIT (Exon-Intron Transformer). We have implemented codon tokenization for this model. This has been validated on the human genome with external validation from the chimpanzee genome. Further benchmarking has implied that our model is better than the existing models in the above tasks.

## 1 Introduction

Identification of exons and introns is a fundamental task in genomics, as it directly enables accurate gene annotation and helps in understanding gene structure in eukaryotic organisms. Exons are the coding regions that are translated to proteins, while introns are the non-coding segments removed during RNA splicing and hence distinguishing between them is essential for reconstruction of functional transcripts and for prediction of protein products. Exon-intron boundary detection plays a crucial role in alternative splicing analysis and identification of regulatory elements that influence gene expressions. Errors in splicing are associated with numerous genetic diseases, making intron-exon prediction for biomedical research and clinical diagnosis.

Deep learning has significantly advanced our capacity to model genomic sequences and identify regulatory elements, such as intron–exon boundaries. Initial methodologies utilized Convolutional Neural Networks (CNNs) [1] and Recurrent Neural Networks (RNNs), including Long Short-Term Memory (LSTM) architectures [2], for tasks ranging from splice site detection to gene structure prediction. However, these architectures encounter significant structural bottlenecks when processing DNA. CNNs are fundamentally constrained by their receptive fields, requiring an impractical depth of layers to capture distal interactions. Similarly, despite their gating mechanisms, RNNs and LSTMs often fail to retain information over the thousands of nucleotides required to model long-range dependencies effectively. This spatial limitation is critical in genomics, where functional regulation frequently depends on non-local context that these legacy models cannot efficiently bridge. Models such as spliceAI [3] demonstrated that deep neural networks trained on large precursor mRNA (pre-mRNA) sequence datasets can accurately predict splice junctions by capturing long range dependencies in DNA sequences. Other models such as SpliceVault [4] work on RNA-sequencing data. However, RNA-sequencing data already represent the expressed and spliced transcripts implying that the intronic regions are largely absent because they are removed during RNA processing. This makes RNA-seq data inherently more biased towards exons and limits its utility for classification of exon-intron boundaries. Although pre-mRNA contains intronic sequences, it is difficult to obtain because it is rapidly processed into mature mRNA and exists only as a transient intermediate. Its instability and short-lived nature restrict the practical application of pre-mRNA data for this purpose.

In recent times, transformer-based architecture has been applied to genomic data, treating DNA sequences as a language. Models such as DNABERT [5] are pretrained on large DNA sequences and have a strong performance for various genomic tasks. Other models such as Nucleotide Transformer [6] further extend this approach by training on massive multi-species datasets, enabling the useful and transferable genomic representations. However, these models use k-mer tokenization, which considers DNA as an overlapping set of 3 or 6 nucleotides. Moreover, these models are typically computationally expensive and require substantial computational resources.We propose a lightweight, transformer-based encoder architecture specifically designed for exon–intron discrimination, splice site recognition, and coding sequence (CDS) classification. Moving away from standard k-mer tokenization, our model utilizes codon-level tokenization to align directly with the biological triplet structure of the genetic code. This approach not only captures essential evolutionary and functional patterns but also maintains a compact vocabulary, significantly reducing computational overhead. By integrating biologically informed preprocessing with an efficient attention mechanism, our model achieves competitive performance in resource-constrained environments. We demonstrate the model’s robustness and cross-species generalizability by training on the human genome and performing external validation on the chimpanzee genome.

## 2 Materials and Methods

### 2.1 Data Sources and Preprocessing

All genomic sequences used in this study were obtained from the human reference genome assembly GRCh38 (Genome Reference Consortium Human Build 38, also known as hg38). The genome sequence was obtained as a multi-chromosome FASTA file (GRCh38.fa), with each chromosome represented as a separate sequence. Gene structure annotations were sourced from Ensembl in the standard Gene Transfer Format (GTF). The GTF file encodes the genomic coordinates of all transcripts, exons, coding sequences (CDS), and strand orientation for every annotated gene in the human genome. The FASTA file was processed in a streaming, chromosome-by-chromosome fashion to avoid loading the entire multi-gigabyte genome into RAM, making the pipeline fully reproducible on standard desktop hardware without requiring a GPU or high-memory server.

#### 2.1.1 Sequence Extraction and Labelling

Labelled sequences for each classification task were extracted directly from the GRCh38 FASTA using coordinates from the Ensembl GTF. For each annotated transcript, exon coordinates were retrieved from the GTF and the corresponding nucleotide sequences were extracted from the FASTA. Six distinct sequence classes were extracted and stored as separate FASTA files.

#### 2.1.2 Task Definitions

The model was trained and evaluated on three hierarchical tasks designed to benchmark its understanding of genomic architecture and have been mentioned in the below table (Table 1):

**Table 1:** Genomic Sequence Classification and Extraction Parameters.

| Sequence Class | Extraction Method | Label in Task |
| --- | --- | --- |
| Exons | Direct coordinate extraction from GTF exon records | Task A: label = 1 |
| Introns | Inferred as inter-exon gaps between consecutive sorted exon boundaries within each transcript | Task A: label = 0 |
| CDS | Direct coordinate extraction from GTF CDS records | Task B: label = 1 |
| Non-CDS | Exonic regions outside CDS boundaries (5'/3' UTRs and non-coding exons) | Task B: label = 0 |
| Splice Donors | 200 nt windows centered on 5' splice junctions (100 nt each side) | Task C: label = 1 |
| Splice Acceptors | 200 nt windows centered on 3' splice junctions (100 nt each side) | Task C: label = 2 |
| No-Splice (negative) | Intronic sequences used as background negative class for Task C | Task C: label = 0 |

**Task A:** Transcript Segmentation (Exon vs. Intron) This task measures the model’s capacity to recognize the structural boundaries of a transcript. Because introns were not explicitly annotated in the GTF, they were inferred computationally as the genomic intervals between consecutive exons within the same transcript.

**Task B:** Coding Potential (CDS vs. Non-CDS) Task B focuses on functional annotation by requiring the model to distinguish between protein-coding sequences (CDS) and non-coding exonic regions, such as Untranslated Regions (UTRs). This evaluates the model’s sensitivity to open reading frames (ORFs) and codon usage bias.

**Task C:** Splice Junction Identification This multi-class task targets the precise recognition of 5’ (Donor) and 3’(Acceptor) splice sites. By centering 200 NT windows on annotated junctions, the model is forced to learn the sequence motifs such as the canonical GT-AG dinucleotides and the surrounding genomic context required for accurate splicing.

#### 2.1.3 Sequence Pre-processing and Refinement

**Introns** were not directly annotated in the GTF and were instead inferred computationally as the genomic intervals between consecutive sorted exon boundaries within the same transcript. **Splice site windows** were extracted as fixed 200 nt windows centered on each annotated splice junction (100 nt upstream and 100 nt downstream). This window captures the canonical GT donor and AG acceptor dinucleotides as complete codon-aligned tokens in the downstream tokenization step.

**Strand orientation:** all sequences from minus-strand features were reverse-complemented to produce canonical 5_′ →_ 3_′_ orientation prior to any further processing.

**Length filtering:** sequences shorter than 50 bp were discarded as insufficient for reliable codon tokenisation. Sequences longer than 5,000 bp were center-cropped to 5,000 bp, retaining the biologically most informative central region.

**IUPAC ambiguity resolution:** ambiguous nucleotide codes (R, Y, S, W, K, M, B, D, H, V and N) present in the reference genome were resolved to the most common unambiguous base prior to tokenization (e.g. N → A, R → A, Y → C).

#### 2.1.4 Codon-Based Tokenization

Nucleotide sequences were converted to integer token sequences using a **codon-based (non-overlapping 3-mer) tokenization scheme**. All 64 possible trinucleotides were enumerated in sorted lexicographic order and assigned integer indices 3–66, with three special tokens reserved: <PAD> = 0, <UNK> = 1, and <MASK> = 2. The resulting vocabulary contains exactly 67 tokens. Tokenization proceeds by reading the sequence in non-overlapping 3-nucleotide windows from the 5_′_ end. This approach was chosen over character-level or overlapping k-mer tokenization for two reasons. First, codons are the fundamental unit of the genetic code, making them the most biologically meaningful sub-sequence unit for distinguishing coding from non-coding regions. Second, a compact vocabulary of 67 tokens avoids embedding matrix sparsity while remaining large enough to capture codon-usage bias, which differs substantially between exonic and intronic regions. Each tokenized sequence was padded to a fixed length using <PAD> tokens (index 0). Padding positions are excluded from the self-attention computation via an attention mask.

### 2.2 Model Architecture

The proposed model uses a Transformer encoder architecture [7] to learn representations from DNA sequences. The model processes codon-tokenized DNA sequences and predicts genomic labels using task-specification classification heads. The architecture consists of an embedding layer, positional encoding, a stack of transformer encoder layers and a classification head. The encoder is shared across tasks so that the model can learn general DNA sequence representations, while different classifier heads are used for different prediction tasks (Fig.1).

**Fig 1.**
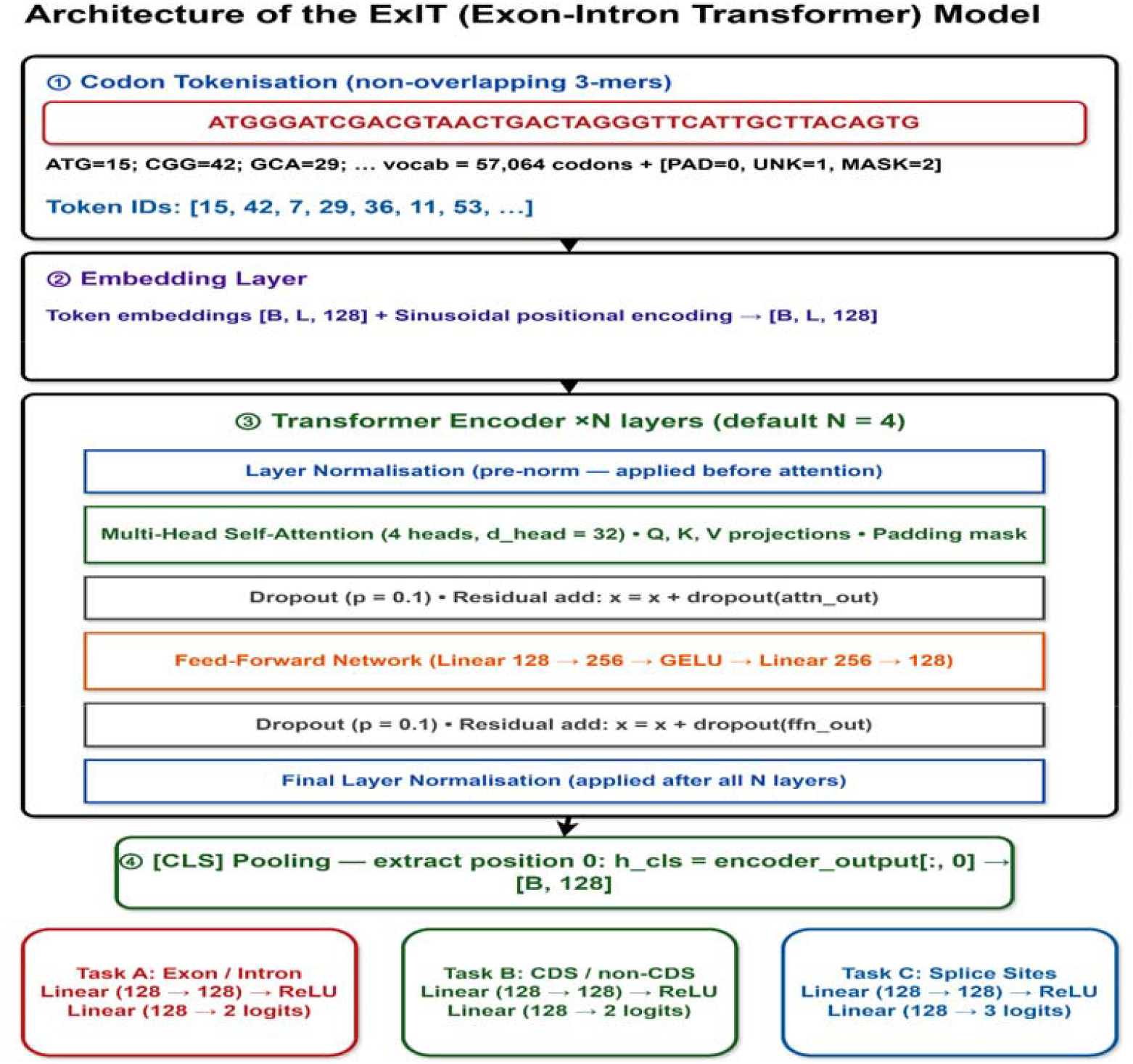
The flow diagram shows the detailed architecture of ExIT

#### 2.2.1 Input Representation

DNA sequences are first converted into a codon-based representation. A codon is a group of 3 nucleotides. We have included all the 64 tokens corresponding to the 64 codons present in the genetic code, plus three special tokens, making the vocabulary size 67. The three additional tokens are used in preprocessing and training pipeline. **PAD token is** used to pad sequences to a fixed length for batch processing while being ignored during attention through padding masks. **UNK token** represents codons that cannot be mapped to the predefined vocabulary, allowing the model to handle ambiguous or corrupted sequence tokens. **MASK token** is used during masked sequence modeling to replace selected tokens so the model learns contextual representations by predicting the masked codons.

Let a DNA sequence be represented as a sequence of codon tokens:

X=(x_1_,x_2_,x_3_,…,x_L_) where L denotes the number of codon tokens in the sequence and each token x_i_ belongs to the vocabulary V. The vocabulary size is defined as |V|=64+3=67 where the additional three tokens correspond to the PAD, UNK, and MASK symbols.

Each token is mapped to a dense vector using an embedding layer. The embedding layer converts token IDs into continuous vector representations of dimension (d_model_). These embeddings allow the model to represent similarities between different codons in a numerical form.

Each token is mapped to a continuous vector representation through an embedding matrix E € R^|V|×dmodel^ where d_model_ represents the embedding dimension. The embedding of token x_i_ is given by e_i_=E[x_i_]. Thus, the embedded representation of the sequence becomes E(X)=(e_1_,e_2_,e_3_,…,e_L_).

A special classification token [CLS] is added at the beginning of each sequence to obtain the global representation of the sequence. This token serves as the summary representation of the entire sequence, allowing the model to aggregate all the contextual information during the self-attention process.

The CLS token is represented by a learnable embedding vector:

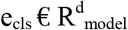

which is trained jointly with the rest of the model parameters. After inserting this token at the beginning of the sequence, the embedded input representation becomes

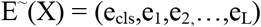

where e_i_ represents the embedding of the i^th^ codon token and L denotes the original sequence length. As a result, the sequence length increases by one position to accommodate the CLS token.

#### 2.2.2 Positional Encoding

Transformers do not inherently capture the order of tokens in a sequence. However the order of codons in a DNA sequence is important in a sequence. For example, the ATG token typically represents the start codon of a sequence however in between the sequence the ATG token typically encodes for methionine amino acid as part of an internal coding region.

Without positional encoding, the transformer would treat both occurrences of **ATG** as identical tokens: *Embedding* (ATG) regardless of where they appear in the sequence. By incorporating positional encoding, the model distinguishes between tokens occurring at different positions:

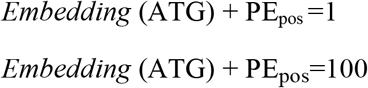

Thus, positional encoding allows the model to differentiate between the same codon appearing at different locations in the sequence and capture biologically meaningful positional patterns in genomic data.

The model uses sinusoidal positional encoding, which is added to the input embeddings. A positional encoding matrix 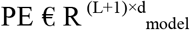 is computed and added to the embeddings. The positional encoding for a position pos and dimension i is defined as:

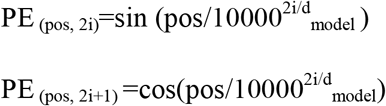

#### 2.2.3 Encoder Architecture

The DNA sequence is learned using a stack of four transformers encoder layers. The encoder processes the input representation obtained after embedding, positional encoding and CLS insertion. Let 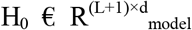 denote the input to the encoder, where L represents the number of codon tokens and d_model_ is the embedding dimension. Each encoder layer consists of two main components:

Multi-head self-attention and a position-wise feed-forward network (FFN). A pre-layer normalization configuration is used to improve stability.

##### 2.2.3.1 Multi-Head Self Attention

The self-attention mechanism allows each token in the sequence to attend to all the other sequences and capture contextual dependencies across the DNA sequence. For an input representation H, query (Q), key(K) and value(V) matrices are computed as

Q= HW_Q_, K=HW_K_, V=HW_V_, where W_Q_, W_K_, W_V_ are learnable projection matrices. The attention process is defined as:

Attention (Q,K,V) = softmax 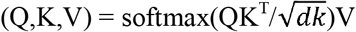V where d_k_ is the dimensionality of the vectors. To improve representation capacity, the model employs multiple self-attention operations which are performed in parallel and concatenated.

MultiHead(Q,K,V) = Concat(head_1_,…,head_h_) W_O_ where h denotes the number of attention heads.

##### 2.2.3.2 Feed Forward Network

Following the attention mechanism, each token representation is processed by a position-wise feed forward network. The FFN consists of two linear transformations with a GELU activation function.

FFN (h) = W_2_(GELU (W_1_h+b_1_)) +b_2_where W_1_ and W_2_ are learnable weight matrices.

##### 2.2.3.3 Activation Function

The feed-forward network employs the Gaussian Error Linear Unit (GELU) as the activation function. GELU is commonly used in transformer architectures because it provides smoother activation compared to traditional functions as ReLU, enabling improved model performance.

In practice, GELU can be approximated as

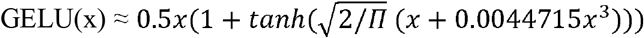

This activation function allows the model to retain small input values while suppressing negative values in a smooth and probabilistic manner, which helps improve the representation learning capability of transformer models. Within the feed-forward network, the GELU activation is applied between the two linear transformations, enabling the model to learn non-linear relationships within the sequence representations.

##### 2.2.3.4 Residual Connections and Layer Normalization

To stabilize training and facilitate efficient gradient propagation in transformer networks, residual connections and layer normalization are applied within each encoder layer. Residual connections allow the model to preserve information from earlier layers without learning additional contextual representations thereby reducing the vanishing gradient problem. In the proposed architecture, a pre-layer normalization is employed. In this mechanism, the input to both the multi-head attention block and feed-forward network is first normalized before applying the corresponding transformation.

Let x denote the input representation to a transformer encoder layer. The computation proceeds as follows.

First, layer normalization is applied before the attention operation:

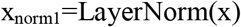

The multi-head self-attention output is then added to the original input through a residual connection:

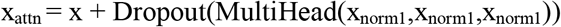

Next, the intermediate representation is normalized again before passing through the feed-forward network:

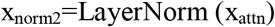

Finally, the output of the feed-forward network is combined with the previous representation using another residual connection:

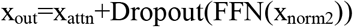

These residual pathways allow the model to retain the original sequence representation while progressively refining contextual information across encoder layers. The use of layer-normalization prior to each transformation improves training stability and enables efficient learning in the model architecture.

##### 2.2.3.5 Adaptive Classifier Head

Following the transformer encoder’s processing of the input sequence, the final hidden state of the [CLS] token is extracted to serve as a high-dimensional summary of the entire genomic sequence. This pooled representation is fed into an adaptive classification head, which consists of a multi-layer perceptron (MLP) with two linear layers separated by a non-linear activation function. While the first layer transforms the encoder’s output within the hidden space to refine feature extraction, the final layer maps these features to the specific dimensionality required by the target task. In this study, our primary focus is the binary discrimination of exons versus introns. However, the architectural flexibility of this head allows the same encoder to be seamlessly adapted for related genomic tasks, including coding sequence (CDS) identification and splice site recognition.

### 2.3 Training Setup

#### 2.3.1 Train / Validation / Test Split

A chromosome-based split strategy was employed throughout this study which has been discussed in Table 2. The genome was partitioned by chromosomes, ensuring that no genomic region appeared in more than one partition. This is the strictest possible data split for human genome data: because chromosomes are non-overlapping by definition, there is zero sequence overlap between training, validation, and test sets. This eliminates all forms of sequence-level data leakage and provides a conservative, unbiased estimate of model generalization.

**Table 2:**
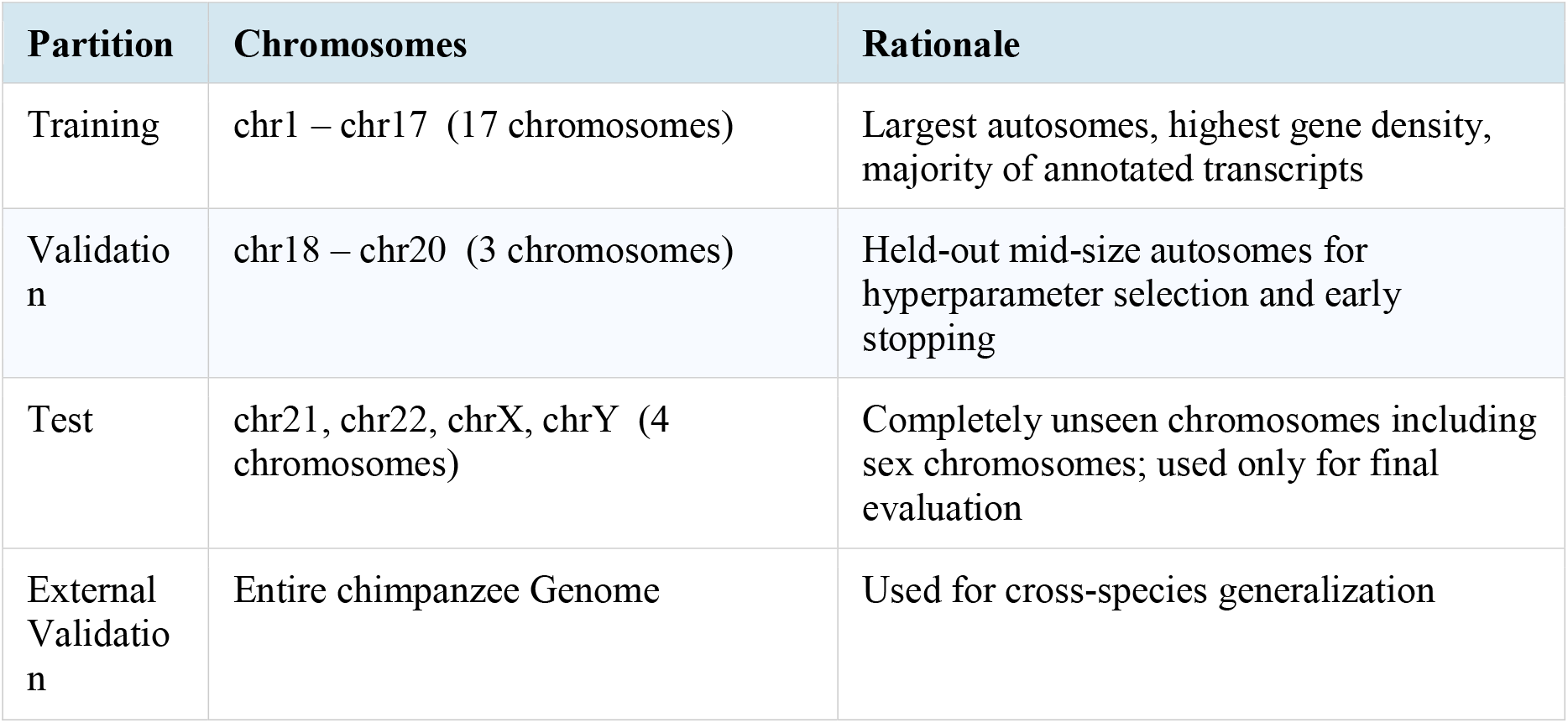
Details of Train/Validation/Test Split.

All primary results reported in this paper use the chromosome-based split. No sequence from any test chromosome was used at any stage of model training or hyperparameter selection. **Importantly, the split is applied at the chromosome level and not at the gene or transcript level**, so all transcripts of a given gene are assigned to the same partition whenever they reside on the same chromosome, further reducing the risk of cross-split gene family contamination.

#### 2.3.2 Encoding to PyTorch Tensors

Tokenized sequences were encoded to PyTorch.pt tensor files for efficient loading during training. Each file stores two tensors: input_ids (shape: [N, T], dtype: int64) and labels (shape: [N], dtype: int64), where N is the number of sequences and T is the number of tokens per sequence.

To accommodate machines with limited RAM, sequences were processed in memory-safe chunks of 10,000 at a time. **Class imbalance** was handled at training time via inverse-frequency class weights passed to the cross-entropy loss function, computed from the training split only. For Task C (splice site detection), where splice site sequences represent a small minority of the available sequences, the intronic negative class was additionally capped at **2 × the number of positive splice sites** during encoding to prevent the model from collapsing to majority-class prediction.

#### 2.3.3 Training Sample Cap

Due to computational constraints (CPU-only training on a consumer-grade Intel Core i7 processor), training was performed on a random subsample of **50**,**000 sequences per task** drawn uniformly from the chromosome-split training partition (chromosomes 1–17). Validation was capped at 7,500 sequences (15% of 50,000) from the validation chromosomes (18–20). Test evaluation was performed on the **full unseen test chromosomes** (chr21, chr22, chrX, chrY) without any capping to provide an unbiased estimate of generalisation performance.

All random subsampling used torch.randperm with a fixed seed of 42 to ensure reproducibility.

#### 2.3.4 Dataset Summary

The below table (Table 3) provides a comprehensive overview of the dataset configuration, detailing the chromosome-based partitioning strategy and the 3-mer codon tokenization parameters. It specifies the window sizes and length-filtering constraints applied across tasks to ensure biological consistency during model training. Additionally, the table summarizes the subsampling and class-weighting protocols used to optimize performance and mitigate the impact of genomic class imbalance.

**Table 3.** Dataset configuration summary.

| Parameter | Value |
| --- | --- |
| Reference genome | GRCh38 (hg38), Homo sapiens |
| Annotation source | Ensembl GTF |
| Genome file format | Multi-chromosome FASTA (.fa) |
| Annotation file format | Gene Transfer Format (.gtf) |
| Split strategy | Chromosome-based only |
| Training chromosomes | chr1 – chr17 |
| Validation chromosomes | chr18 – chr20 |
| Test chromosomes | chr21, chr22, chrX, chrY |
| Tokenisation | Codon-based non-overlapping 3-mer, vocab size = 67 |
| Min sequence length | 50 bp (all tasks) |
| Max sequence length | 5,000 bp (centre-cropped) |
| Task A window | 512 nt → 170 tokens (both experiments) |
| Task B window | 512 nt → 170 tokens (Exp1) / 333 nt → 111 tokens (Exp2) |
| Task C window | 200 nt → 66 tokens (both experiments) |
| Max training samples | 50,000 per task (random subsample, seed = 42) |
| Max validation samples | 7,500 per task |
| Test evaluation | Full test chromosomes (no capping) |
| Class imbalance handling | Inverse-frequency class weights in cross-entropy loss |
| Task C negative cap | $2 \times$ number of positive splice sites |
| Tensor format | PyTorch .pt (input_ids: int64, labels: int64) |

### 2.4 Hyperparameter Optimization

#### 2.4.1 Search Strategy

Hyperparameter optimization was performed via **random search** over the space defined in Table 3. A total of **50 randomly sampled configurations** were evaluated per task (150 runs in total across all three tasks). Each configuration was trained for **3 epochs** on a capped subsample of 20,000 training sequences to serve as a fast performance proxy, keeping the total search time within approximately two hours on a consumer-grade Intel Core i7 CPU.

To prevent the search from accidentally discarding the already-working configuration, the **current default hyper parameters were always evaluated as run #1**, with the remaining 49 runs drawn randomly. For each hyperparameter independently, the value that yielded the highest mean MCC across all runs in which it appeared was selected as the best value. This marginal analysis strategy identifies the most impactful value per parameter without requiring exhaustive grid search over all combinations. All the chosen values for each hyperparameter has been mentioned in Table 4

**Table 4.**
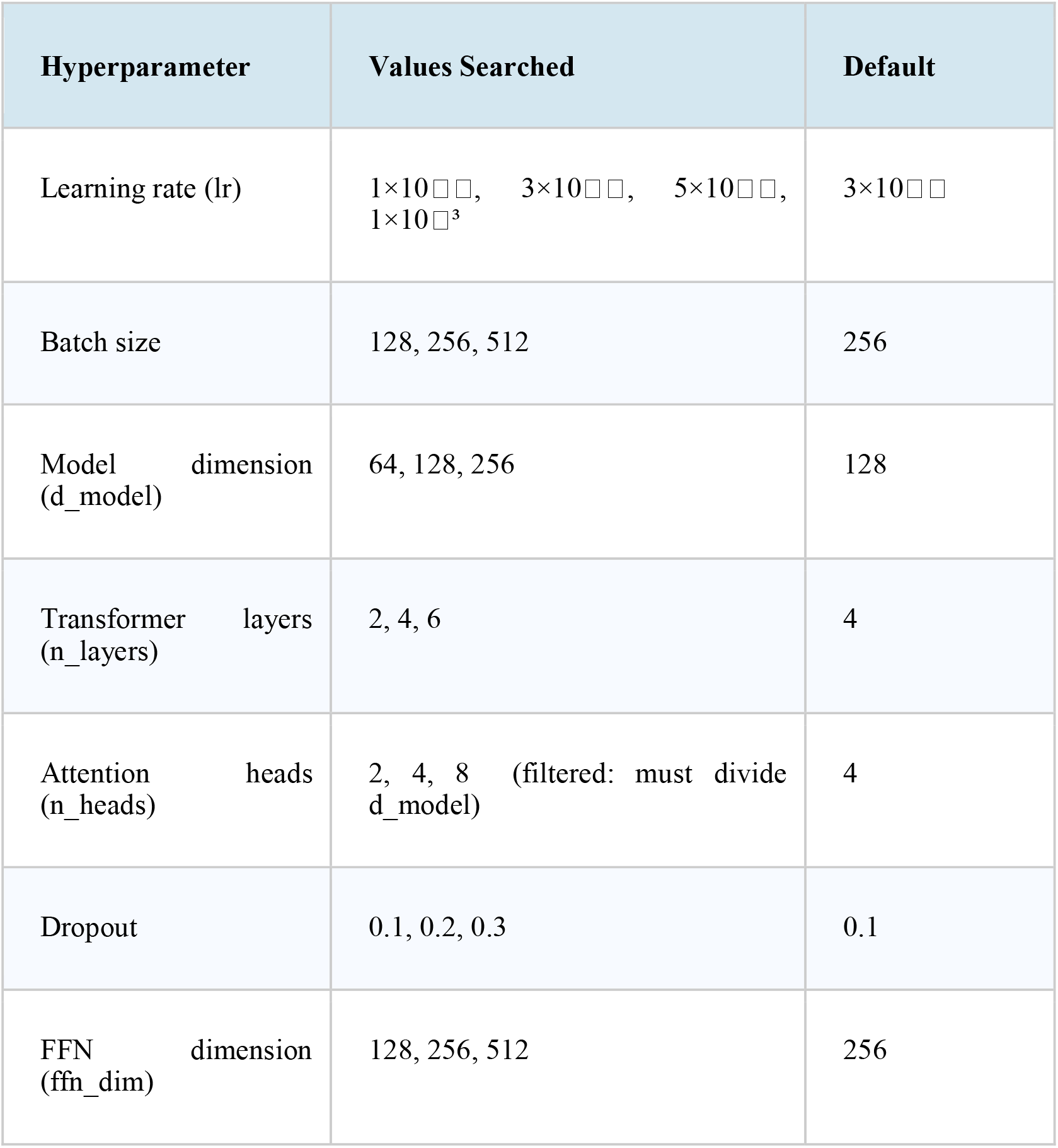
Hyperparameter search space.

#### 2.4.2 Best Hyperparameters Found

Table 5 reports the best hyperparameter values identified by the random search for each task. The search converged on learning rate (**lr) = 0.001** for all three tasks, which is higher than the default of 3×10□□. This suggests that with only 3 search epochs and 20,000 training sequences, a higher learning rate allows faster convergence within the limited budget. The default dropout of 0.1 was retained across all tasks, indicating that the original regularization strength was already appropriate.

**Table 5.**
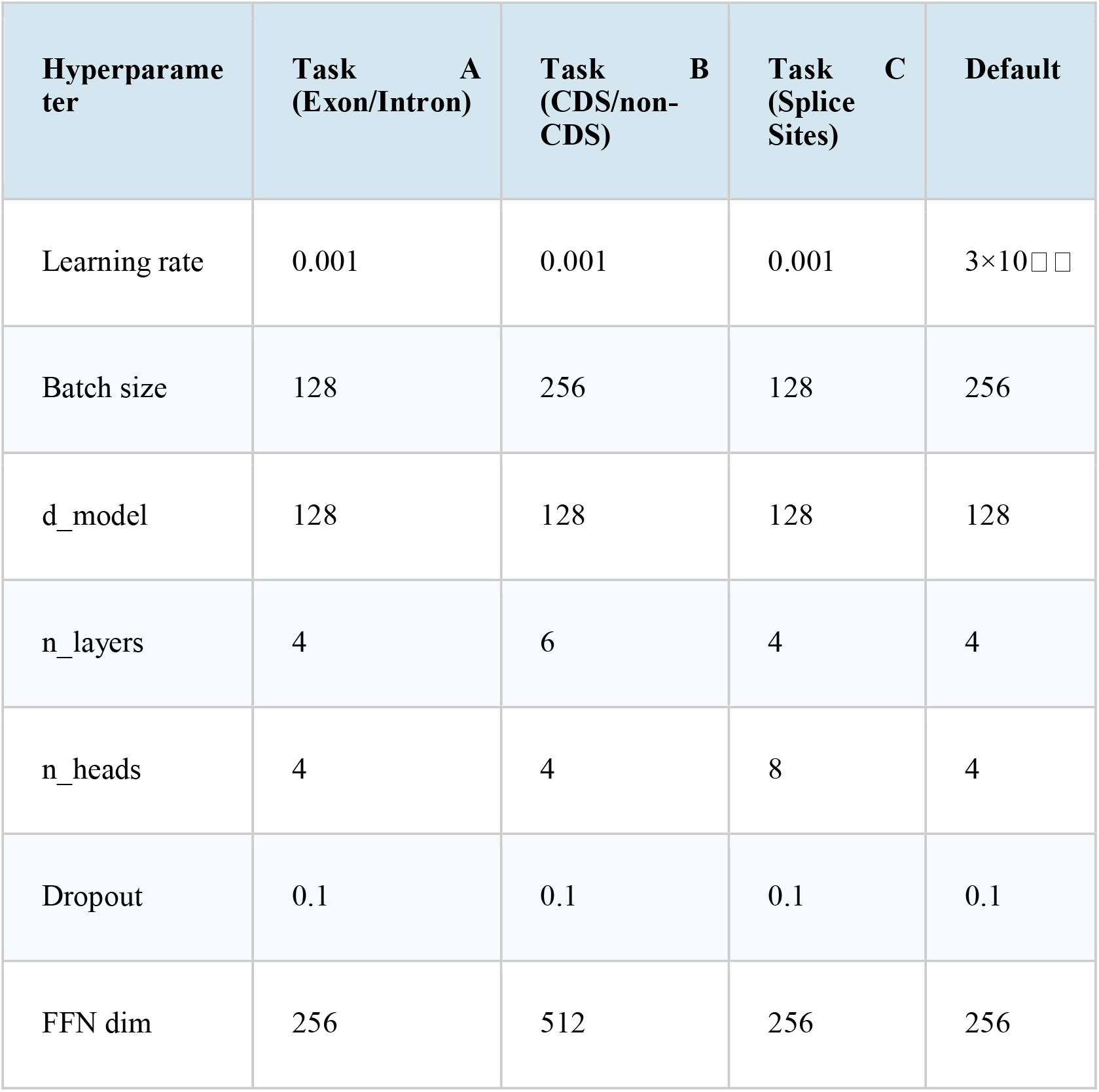
Best hyperparameters per task identified by random search.

Notable task-specific differences emerged from the search. **Task B** benefited from a deeper architecture with **6 transformer layers** compared to 4 for the other tasks, and a wider feed-forward network (FFN dim = 512 vs 256). This is consistent with CDS/non-CDS classification being a more complex task requiring the model to capture longer-range codon usage patterns across the 333 nt window. **Task C** converged on **8 attention heads** rather than 4, which may reflect the need to simultaneously attend to the GT donor and AG acceptor signals at different positions within the 200 nt splice junction window.

### 2.5. Evaluation Metrics

In the development and testing of the ExIT (Exon-Intron Transformer) model, the **Matthews Correlation Coefficient (MCC)** was utilized as the primary performance metric for all genomic classification tasks, including task A (Exon/Intron), task B (CDS/non-CDS), and task C (Splice Sites).

Definition and Mathematical Framework

The MCC is a comprehensive measure of binary classification quality that incorporates all four components of the confusion matrix. It is formally defined by the following equation:

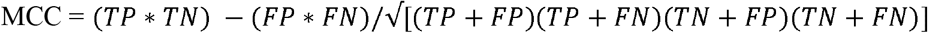

The resulting value ranges from **-1** (total disagreement) to **+1** (perfect prediction), with **0** indicating a performance no better than random chance. The shift from accuracy to MCC as the lead metric was necessitated by the specific nature of genomic data and the architectural goals of the ExIT project:

- **Handling Class Imbalance:** Genomic sequences are inherently imbalanced; for instance, regulatory regions like splice sites are significantly rarer than non-functional sequences. Accuracy can be deceptively high in these scenarios by simply favoring the majority class, whereas MCC remains an informative score even when class sizes differ greatly.
- **Symmetric Evaluation:** Unlike accuracy, which treats all correct predictions equally, MCC yields a high score only if the model performs well in both the positive and negative categories (TP and TN). This is vital for biological applications where missing an actual regulatory region (False Negative) is as critical as a false identification (False Positive).

## 3. Results and Discussion

This section presents results across four subsections: (i) the effect of sequence window length on classification performance (Case Studies 1 and 2), (ii) performance after hyperparameter tuning (the final tuned models), (iii) cross-species generalisation to chimpanzee, and (iv) a quantitative comparison with the Nucleotide Transformer 500M foundation model (NT-500M) under linear probe conditions. All evaluations use the chromosome-based held-out test set (chr21, chr22, chrX, chrY), which was never seen during training or hyperparameter selection.

### 3.1 Effect of Sequence Window Length on Classification Performance

#### 3.1.1 Case Study 1: Uniform 512 nt Window (Experiment 1)

The first experiment (Exp1) applied a uniform 512 nucleotide (nt) window to all three tasks, yielding 170 codon tokens per sequence after non-overlapping codon tokenisation. This single window length served as the baseline configuration and established the extent to which a biologically agnostic preprocessing decision constrains downstream classification performance.

Results across the three tasks were markedly heterogeneous (Table 6). Task A (exon/intron classification) achieved strong performance, with MCC = 0.8719, macro F1 = 0.9343, and accuracy = 0.9343, indicating that a 512 nt window captures sufficient sequence context for reliable exon/intron discrimination. In contrast, Task B (CDS vs non-CDS) performed poorly under the same window, yielding MCC = 0.2541 and accuracy = 0.5892 — only marginally above chance for a binary task. Task C (splice site classification) attained MCC = 0.4972 and accuracy = 0.6535, representing moderate performance.

**Table 6.**
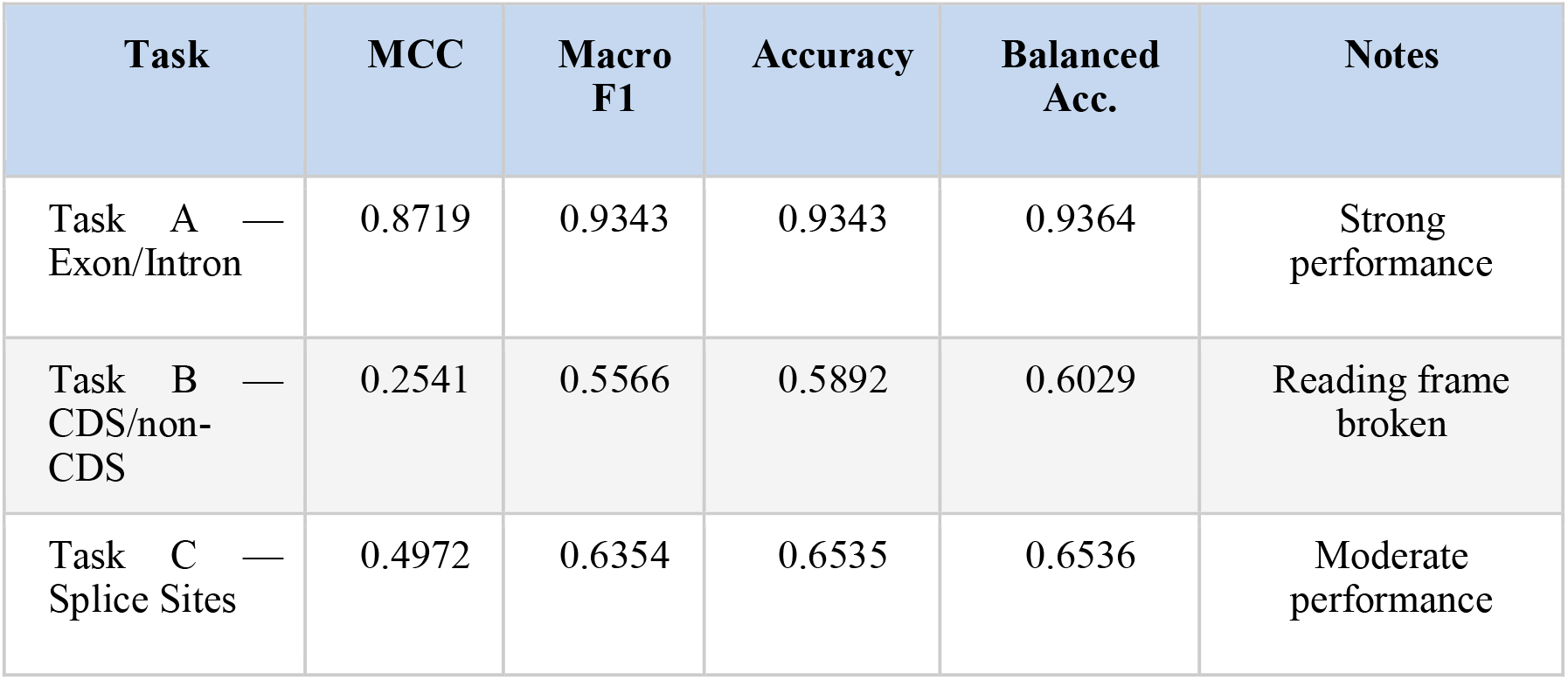
Experiment 1 (Exp1) results — uniform 512 nt window applied to all three tasks.

| Task | MCC | Macro F1 | Accuracy | Balanced Acc. | Notes |
| --- | --- | --- | --- | --- | --- |
| Task A — Exon/Intron | 0.8719 | 0.9343 | 0.9343 | 0.9364 | Strong performance |
| Task B — CDS/non-CDS | 0.2541 | 0.5566 | 0.5892 | 0.6029 | Reading frame broken |
| Task C — Splice Sites | 0.4972 | 0.6354 | 0.6535 | 0.6536 | Moderate performance |

The degraded performance on Task B under the 512 nt window is attributable to a systematic biological mismatch: the uniform window truncates or pads CDS sequences without regard for codon triplet boundaries, thereby disrupting reading frame continuity. Because CDS sequences are defined precisely by in-frame codon structure, any window that misaligns the reading frame introduces artificial stop codons and corrupts the very signal the model must learn. This observation motivated Experiment 2.

#### 3.1.2 Case Study 2: Biologically Appropriate Windows (Experiment 2)

The second experiment (Exp2) replaced the uniform 512 nt window with task-specific window lengths grounded in the biological properties of each sequence type. Task A was left unchanged, as its performance was already excellent. Task B was re-encoded using a 333 nt window (111 codons), chosen to be an exact multiple of three nucleotides so that all extracted CDS sequences begin and end on a codon boundary, preserving reading frame integrity. Task C was re-encoded using a 200 nt window centred on each splice junction (100 nt per side), capturing the canonical GT-AG donor/acceptor dinucleotide signals and their flanking exonic and intronic context.

The effect on Task B was substantial. Correcting the reading frame alignment increased MCC from 0.2541 to 0.4452, a gain of +0.191 MCC points — the single largest performance improvement observed across all experiments in this study, exceeding the gains attributable to hyperparameter tuning. This result provides direct empirical evidence that reading frame preservation is a primary determinant of CDS classification performance when using codon-based tokenisation.

Task C showed a modest decline under the biologically motivated 200 nt window (MCC: 0.4972 →0.4572, Δ = −0.040). Inspection of the per-class confusion matrices revealed that the narrower window degraded the classification of no-splice (intronic background) sequences specifically, as the reduced context made it harder to distinguish background intronic regions from true splice junctions. Experiment 1’s 512 nt window was therefore retained as the superior configuration for Task C. Table 7 summarises the Exp1 versus Exp2 comparison.

**Table 7.**
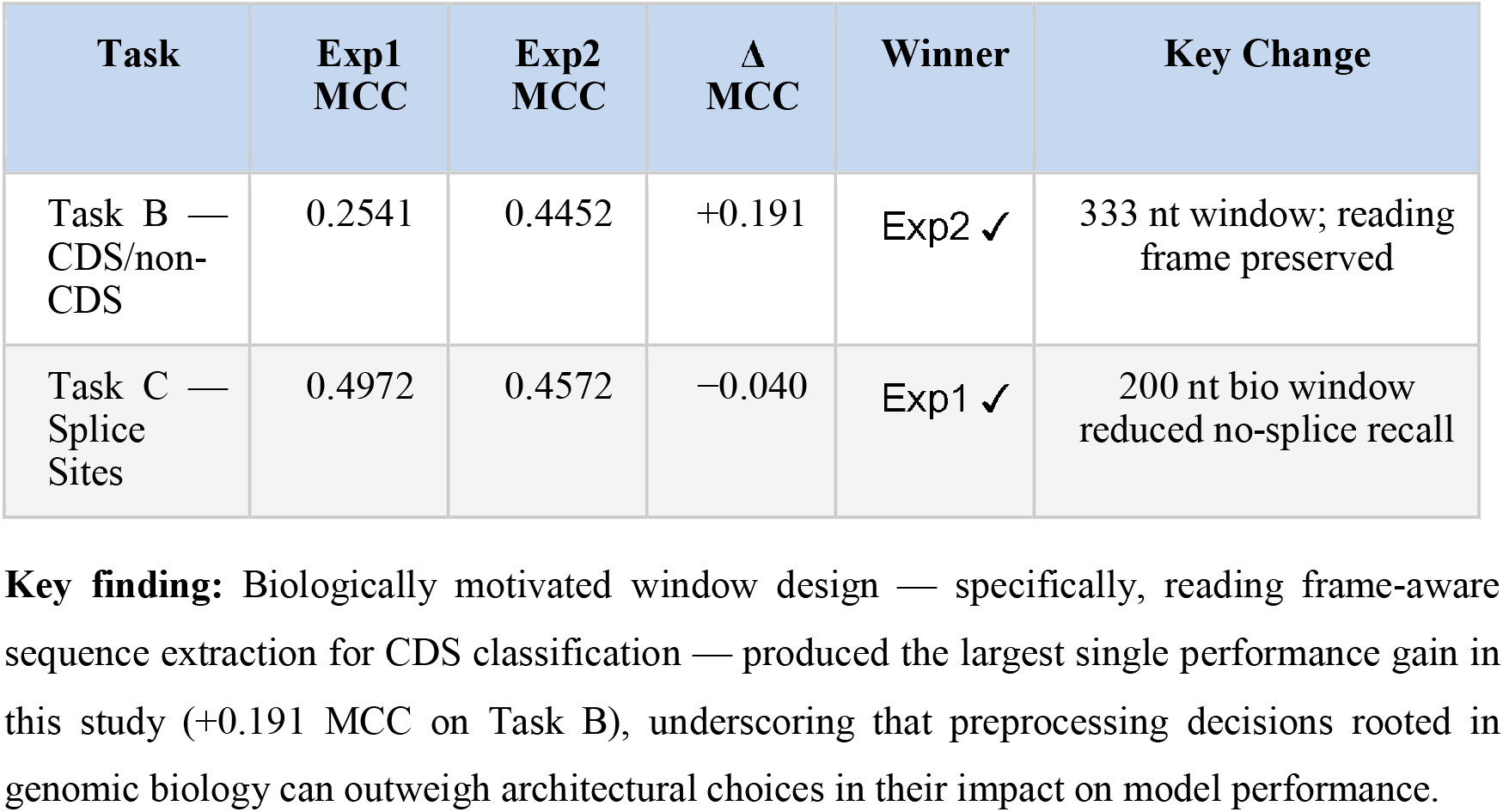
Experiment 1 vs Experiment 2 comparison. Task A unchanged (MCC = 0.877 retained). Best configuration per task highlighted.

| Task | Exp1<br>MCC | Exp2<br>MCC | $\Delta$<br>MCC | Winner | Key Change |
| --- | --- | --- | --- | --- | --- |
| Task B —<br>CDS/non-CDS | 0.2541 | 0.4452 | +0.191 | Exp2 ✓ | 333 nt window; reading<br>frame preserved |
| Task C —<br>Splice<br>Sites | 0.4972 | 0.4572 | -0.040 | Exp1 ✓ | 200 nt bio window<br>reduced no-splice recall |

**Key finding:** Biologically motivated window design — specifically, reading frame-aware sequence extraction for CDS classification — produced the largest single performance gain in this study (+0.191 MCC on Task B), underscoring that preprocessing decisions rooted in genomic biology can outweigh architectural choices in their impact on model performance.

### 3.2 Tuned Model Performance

Following the window-length case studies, a random hyperparameter search was conducted (50 combinations per task, 150 total runs) using a three-epoch proxy training procedure with 20,000-sequence subsets. The best-performing hyperparameter configuration for each task (Table 8) was then used to train fresh models for ten full epochs on the complete training data. These tuned models represent the final configurations used in all downstream evaluations.

**Table 8.**
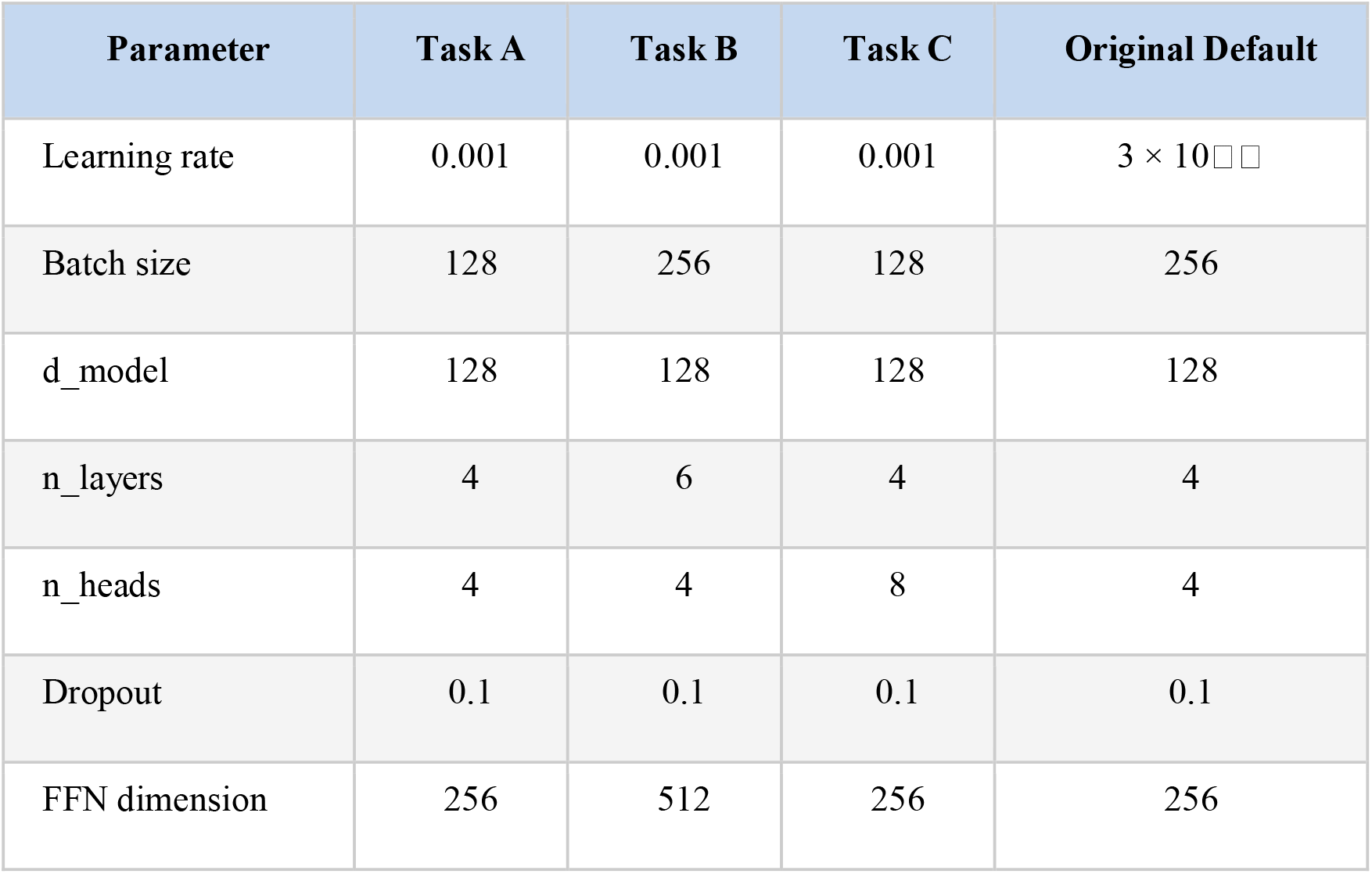
Best hyperparameters identified by random search. All tasks converged on lr = 0.001 (higher than the 3 × 10□□ default), while dropout = 0.1 was consistently optimal.

All three tasks benefited from the higher learning rate (lr = 0.001 vs the 3 × 10□□ default), suggesting that the model’s relatively compact size (∼1.5M parameters) allows faster convergence without overfitting under the three-epoch proxy criterion. Task B required a deeper architecture (6 transformer layers, FFN = 512) compared to Tasks A and C (4 layers, FFN = 256), consistent with the longer 333 nt input window demanding greater representational capacity to capture longer-range codon-usage patterns. Task C uniquely benefited from 8 attention heads (vs the default of 4), plausibly because the model must simultaneously attend to the GT donor dinucleotide and the AG acceptor dinucleotide at opposite ends of the splice junction window.

The tuned models achieved consistent improvements over the best pre-tuning baselines on all three tasks (Table 9). Task A improved from MCC = 0.8719 to MCC = 0.9048 (Δ = +0.033), with accuracy and balanced accuracy both reaching 0.9525–0.9527 across 208,405 test sequences. Task B showed the largest absolute tuning gain: MCC rose from 0.4452 to 0.5557 (Δ = +0.111), with balanced accuracy reaching 0.7806 across 48,162 test sequences. Task C improved modestly from MCC = 0.4972 to MCC = 0.5176 (Δ = +0.020), with accuracy and balanced accuracy of 0.6538.

**Table 9.**
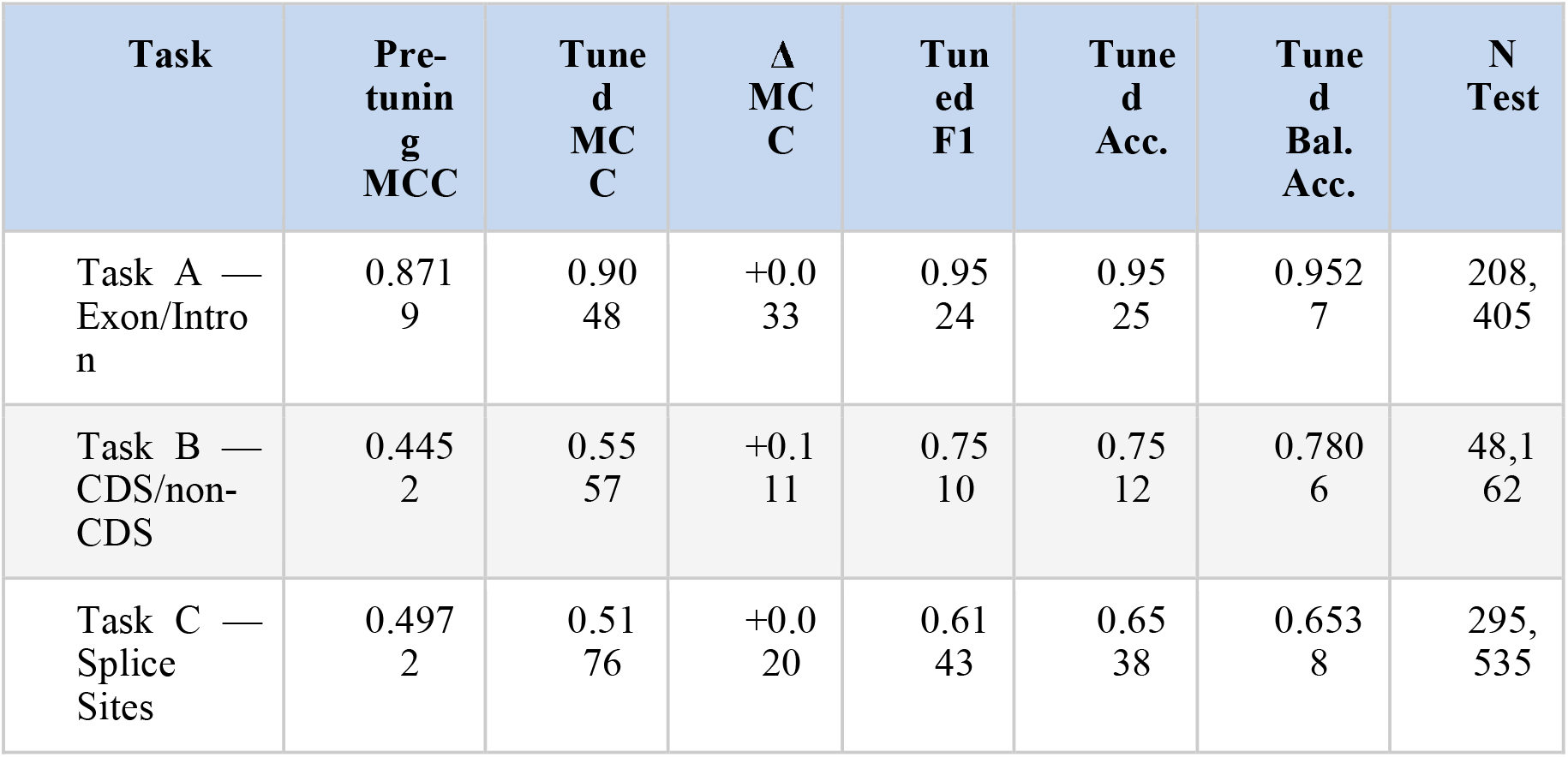
Final tuned model results. Pre-tuning MCC reflects the best result from Exp1/Exp2 for each task.

Despite the overall improvement, Task C retained a systematic classification error: 79,468 acceptor splice site sequences were predicted as donors. The no-splice background class was correctly identified with high recall (97.3%), indicating that the model reliably distinguishes splice sites from intronic background but struggles to differentiate donor from acceptor junction types. This donor/acceptor confusion may stem from the high sequence similarity between the two junction types in their flanking regions, which share the complementary strand sequences of the same splice signals.

### 3.3 Cross-Species Generalisation: Chimpanzee Validation

To evaluate the generalizability of the learned sequence representations, the Task A model (trained exclusively on human GRCh38 chromosomes 1–17) was applied without any retraining or fine-tuning to chimpanzee (Pan troglodytes) exon and intron sequences. A stratified 10% subsample of chimpanzee genomic sequences was drawn (seed = 42), yielding 95,176 sequences (46,439 introns; 48,737 exons) for evaluation.

The model transferred to the chimpanzee genome with only marginal degradation across all metrics (Table 10). MCC declined from 0.8719 to 0.8462 (Δ = −0.0257), retaining 97.1% of human performance. Accuracy and balanced accuracy fell by 1.2% and 1.4% respectively, and the AUC dropped by less than 0.001. Per-class analysis revealed that intron precision was largely preserved (0.9285 → 0.9262), while exon precision declined modestly (0.9401 → 0.9203), and exon recall remained high at 0.9305. The model’s average softmax confidence on exon sequences was 0.931, compared to 0.886 on intron sequences, suggesting that exonic features are more robustly encoded.

**Table 10.**
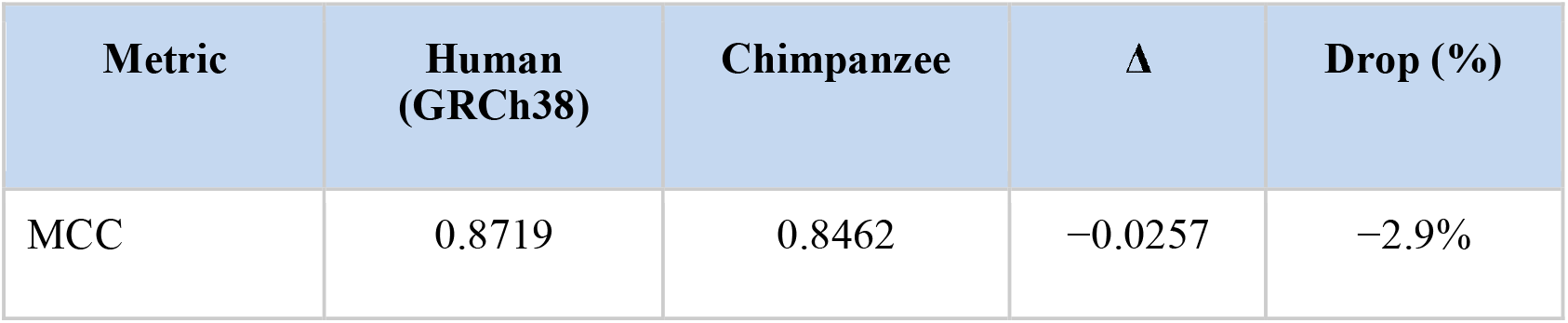
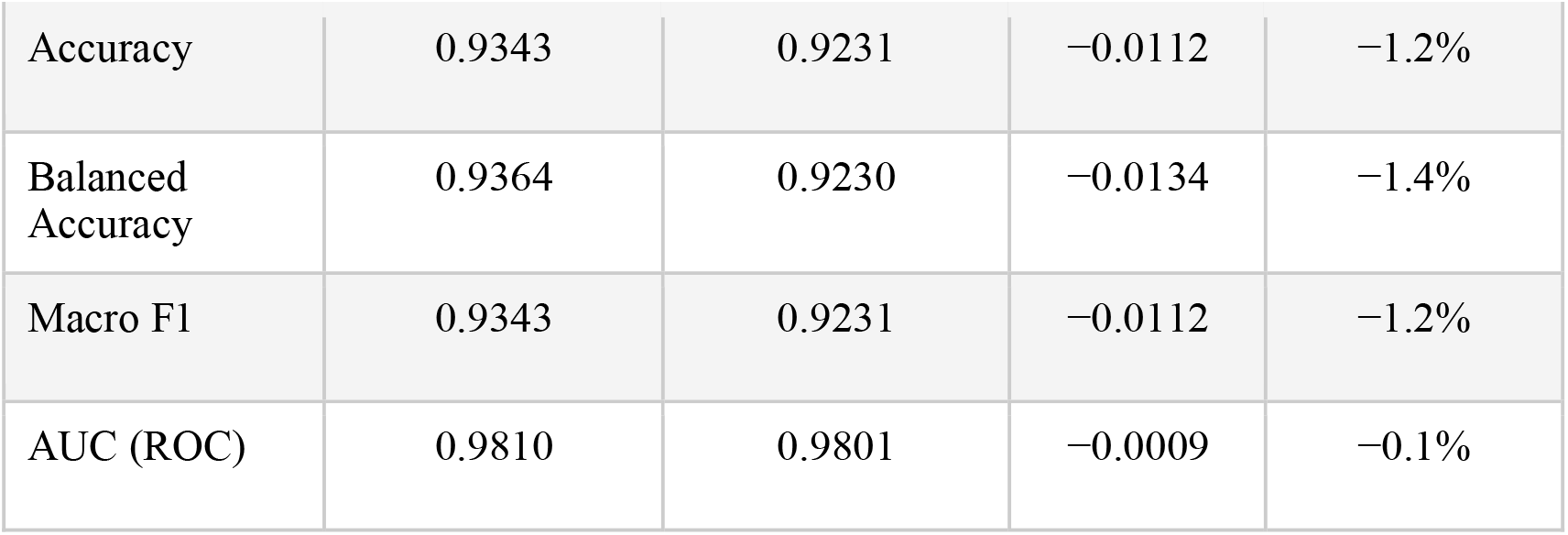
Human vs chimpanzee performance — Task A model applied without retraining. MCC retention: 97.1% of human performance.

Of the 95,176 chimpanzee sequences evaluated, 42,510 introns and 45,351 exons were correctly classified, with 3,929 introns misclassified as exons and 3,386 exons misclassified as introns. The symmetry of these error rates across classes is consistent with the model applying a generalisable decision boundary rather than overfitting to class-specific human genomic biases.

These results provide strong evidence that the learned exon/intron representations reflect deeply conserved sequence features across primate genomes. The near-complete retention of performance (MCC retention: 97.1%) without any chimpanzee data exposure during training suggests that the codon-based transformer has captured biologically fundamental properties of gene structure — such as GC content distributions, splice signal conservation, and exonic compositional biases that are evolutionarily stable across the human-chimpanzee divergence (∼6 million years).

### 3.4 Benchmarking Against the Nucleotide Transformer 500M

To contextualize the performance of the proposed Codon Transformer relative to a large-scale genomic foundation model, we evaluated the Nucleotide Transformer 500M (NT-500M; InstaDeepAI/nucleotide-transformer-500m-human-ref) using the standard linear probe protocol described in the original NT publication (Dalla-Torre et al., Nature Methods, 2024). All 500M NT parameters were frozen; only a single linear layer (1,280 × n_classes weights) was trained on 5,000 sequences per task for five epochs. NT embeddings were obtained by mean-pooling the last hidden layer (embedding dimension: 1,280) across all non-padding positions. Evaluation was performed on the same chromosome-based test set (chr21, chr22, chrX, chrY) used throughout this study.

The Codon Transformer outperformed NT-500M under linear probe conditions on all three tasks (Table 11). On Task A, our Exp1 model achieved MCC = 0.8719 versus NT-500M’s MCC = 0.7758 (Δ = +0.096 in our favour), and the tuned model further extended this advantage by 0.129. The gap was larger on Tasks B and C: our Exp2 model surpassed NT-500M on Task B by 0.149 MCC, and on Task C by 0.290 MCC — the largest margin across all comparisons. Full metrics including F1, balanced accuracy, and accuracy are presented in Table 12.

**Table 11.** MCC comparison — NT-500M linear probe vs Codon Transformer (Exp1/2 best and tuned). Our model outperforms NT-500M on all three tasks despite being ∼333× smaller.

| Model | Params | Task A<br>MCC | Task B<br>MCC | Task C<br>MCC | Training |
| --- | --- | --- | --- | --- | --- |
| NT-500M (linear probe) | 500M | 0.7758 | 0.2965 | 0.2077 | Pretrained (3,202 genomes) |
| Codon Transformer (ours) — Exp1/2 | $\sim 1.5M$ | 0.8719 | 0.4452 | 0.4972 | From scratch |
| Codon Transformer (ours) — Tuned | $\sim 1.5M$ | 0.9048 | 0.5557 | 0.5176 | From scratch |

**Table 12.** Full metric comparison — NT-500M linear probe vs Codon Transformer (Exp1/Exp2 best configurations). NT-500M embedding dimension: 1,280. Codon Transformer probe: not applicable (trained end-to-end).

| Task | Model | MCC | Macro F1 | Balanced Acc. | Accuracy |
| --- | --- | --- | --- | --- | --- |
| Task A | NT-500M (linear probe) | 0.7758 | 0.8878 | 0.8883 | 0.8880 |
| Task A | Codon Transformer (ours) | 0.8719 | 0.9343 | 0.9364 | 0.9343 |
| Task B | NT-500M (linear probe) | 0.2965 | 0.6283 | 0.6415 | 0.6343 |
| Task B | Codon Transformer (ours) | 0.4452 | 0.5566 | 0.6029 | 0.5892 |
| Task C | NT-500M (linear probe) | 0.2077 | 0.4203 | 0.4630 | 0.4629 |
| Task C | Codon Transformer (ours) | 0.4972 | 0.6354 | 0.6536 | 0.6535 |

The per-class breakdown for NT-500M reveals specific weaknesses that explain the lower aggregate MCC scores. On Task C, NT-500M achieved donor recall of only 0.116 (11.6%), meaning fewer than one in eight true donor splice sites were correctly identified, despite reasonable acceptor recall (0.557). On Task B, NT-500M exhibited a strong bias towards the CDS class (CDS recall = 0.799 vs non-CDS recall = 0.485), suggesting that the NT pretraining distribution — which emphasizes protein-coding sequence — may bias the linear probe towards CDS predictions regardless of true class. On Task A, NT-500M performed competitively at the class level, with intron recall of 0.893 and exon recall of 0.883, yet still fell 0.096 MCC points below our model.

These results should not be interpreted as a criticism of NT-500M, which was not designed for fine-grained task-specific classification under linear probe conditions. Rather, the comparison demonstrates that a purpose-built, lightweight transformer (∼1.5M parameters) with biologically motivated tokenisation and task-appropriate preprocessing achieves stronger performance on these three genomic classification tasks than a 333-fold larger foundation model evaluated under the standard linear probe protocol. This finding supports the broader argument that domain-specific inductive biases — here, codon-aligned tokenisation and reading frame-aware sequence extraction — can be more informative than sheer model scale for targeted genomic classification problems.

## 4. Discussion

The experimental results of the ExIT (Exon-Intron Transformer) model demonstrate that high-performance genomic modeling is achievable through a lightweight architecture when combined with biologically informed design. A central contribution of this study is the implementation of codon-level tokenization, which directly incorporates the biological triplet structure of the genetic code into the model’s architecture. By aligning the input vocabulary with these fundamental units, ExIT is inherently designed to capture functional and evolutionary patterns encoded in reading frames.

This alignment proved essential in the transition from Case Study 1 to Case Study 2. In Case Study 1, the use of a uniform 512 nt window resulted in a poor MCC of 0.254 for CDS classification, largely because arbitrary cropping disrupted the triplet sequence. However, by introducing a 333 nt window specifically chosen for its exact divisibility by three, the tuned model’s performance for Task B surged to 0.5557. This suggests that architectural alignment with molecular biology and the preservation of the reading frame are as vital as the neural network depth itself.

Furthermore, the model’s robust MCC of 0.9048 for Exon-Intron classification (Task A) confirms that a 1.5M parameter model can compete with vastly larger state-of-the-art architectures. When compared to the Nucleotide Transformer (NT-500M), which utilizes 500M parameters, ExIT offers a massive 333x reduction in computational overhead without sacrificing the ability to identify core regulatory regions. The practical utility of ExIT is further underscored by its training efficiency and optimized data-handling pipeline. Developed and validated within a constrained 8GB RAM, CPU-only environment, the model utilizes a strategic sample-capping mechanism (50,000 sequences) for both training and validation sets to ensure rapid RAM processing and prevent memory overflow. To further manage resource consumption, the training architecture employs a sequential chromosome loading strategy, processing one chromosome at a time. This approach proves that sophisticated transformer performance is accessible to researchers without high-end GPU clusters or massive memory overhead. Finally, the successful external validation on the chimpanzee genome provides strong evidence of the model’s generalizability. By learning fundamental codon-usage biases and splice-site motifs rather than species-specific human sequences, ExIT represents a scalable, resource-efficient tool for cross-species genomic annotation and the study of evolutionary conserved regulatory elements.

Beyond its architectural efficiency, the **Exon-Intron Transformer (ExIT)** offers significant utility in democratizing genomic research. Its lightweight design enables the annotation of non-model organisms on standard workstation hardware, bypassing the need for high-performance computing (HPC) infrastructure. Furthermore, ExIT’s performance across diverse taxa establishes it as a robust tool for comparative genomics and evolutionary analysis.

In synthetic biology, the model serves as a sequence validator by applying its learned genomic syntax to verify the structural integrity of engineered DNA. Finally, its minimal computational footprint facilitates real-time, decentralized inference in field-based or resource-constrained environments, effectively bridging the gap between raw data acquisition and functional insight.

## Data availability

The datasets supporting the findings of this study are available in public repositories as detailed below:

- Human (Homo sapiens) Genome Sequence (FASTA) Data: The genomic sequence was derived from the human reference genome assembly GRCh38 (Primary Assembly) via the Ensembl database. Link : Index of /pub/release-109/fasta/homo_sapiens/dna.
- Gene Annotations (GTF) of Homo sapiens: Genomic annotations were obtained from the Ensembl Release 109 GTF file. Link : Index of /pub/release-109/gtf/homo_sapiens
- Chimpanzee (Pan troglodytes) Genome Sequence (FASTA) Data: The genomic sequence was derived from the chimpanzee reference genome assembly Pan_tro_3.0 (Toplevel) via the Ensembl database. Link : Index of /pub/release-111/fasta/pan_troglodytes/dna
- Gene Annotations (GTF) of chimpanzee: Genomic annotations were obtained from the Ensembl Release 111 GTF file. Link : Index of /pub/release-111/gtf/pan_troglodytes

## Code availability

The code used to develop the ExIT model are available via GitHub int this particular repository: pavanKarthik2006/ExIT-Exon-Intron-Transformer-

## Acknowledgements

A.B.D. thanks the National Institute of Technology, Warangal, for providing a computational facility.

## Contributions

A.S.P.K. performed the experiment. A.B.D. conceived and planned the experiments.

A.S.P.K. and A.B.D. wrote the manuscript. All authors discussed the results and contributed to the final manuscript.

## References

[1] F. Ben Nasr Barber and A. Elloumi Oueslati, “Human exons and introns classification using pre-trained Resnet-50 and GoogleNet models and 13-layers CNN model,” Journal of Genetic Engineering and Biotechnology, vol. 22, no. 1, p. 100359, Mar. 2024, doi: 10.1016/j.jgeb.2024.100359.

[2] B. Lee, T. Lee, B. Na, and S. Yoon, “DNA-Level Splice Junction Prediction using Deep Recurrent Neural Networks,” Dec. 16, 2015, arXiv: arXiv:1512.05135. doi: 10.48550/arXiv.1512.05135.

[3] B. A. Jónsson et al., “Transformers significantly improve splice site prediction,” Commun Biol, vol. 7, no. 1, p. 1616, Dec. 2024, doi: 10.1038/s42003-024-07298-9.

[4] R. Dawes et al., “SpliceVault predicts the precise nature of variant-associated mis-splicing,” Nat Genet, vol. 55, no. 2, pp. 324–332, Feb. 2023, doi: 10.1038/s41588-022-01293-8.

[5] Y. Ji, Z. Zhou, H. Liu, and R. V. Davuluri, “DNABERT: pre-trained Bidirectional Encoder Representations from Transformers model for DNA-language in genome,” Bioinformatics, vol. 37, no. 15, pp. 2112–2120, Aug. 2021, doi: 10.1093/bioinformatics/btab083.

[6] H. Dalla-Torre et al., “Nucleotide Transformer: building and evaluating robust foundation models for human genomics,” Nat Methods, vol. 22, no. 2, pp. 287–297, Feb. 2025, doi: 10.1038/s41592-024-02523-z.

[7] A. Vaswani et al., “Attention Is All You Need,” Aug. 02, 2023, arXiv: arXiv:1706.03762. doi: 10.48550/arXiv.1706.03762.

